# A high-throughput immunopeptidome platform for MHC II alleles to characterize antigen-specific CD4^+^ T cells

**DOI:** 10.1101/2025.06.05.658000

**Authors:** Jing Chen, Xu Zhu, Jun Huo, Shang Wu, Ting Zhou, Chunyu Cheng, Hao Dong, Yan Li, YuXin Chen, Xianchi Dong

## Abstract

CD4^+^ T cells play a pivotal role in adaptive immunity, recognizing peptide antigens presented by MHC II molecules during infections and tumor development. Identifying immunodominant MHC II epitopess is essential for understanding CD4^+^ T cell responses; however, current methods such as mass spectrometry, suffer from low sensitivity and throughput, while computational algorithms show variable accuracy. To overcome these challenges, we developed EliteMHCII, a high-throughput immunopeptidome profiling platform that identifies antigen-derived MHC II epitopes and measures peptide binding affinity across 24 globally common MHC II alleles. Using EliteMHCII, we assessed the immunodominant epitopes of the SARS-CoV-2 RBD protein. Validation in vaccinated individuals and humanized mouse models revealed a strong correlation between high-affinity peptides and robust CD4^+^ T cell responses, while low-affinity peptides failed to elicit responses. Therefore, our immunopeptidome profiling platform, EliteMHCII, serves as a rapid, high throughput, feasible platform for CD4^+^ T cell epitope discovery at a global populational level in the context of infectious diseases and cancer immunotherapy.

## Introduction

CD4^+^ T cells play a crucial role in the adaptive immune system^1,2^, which could orchestrate the generation of high-quality, long-lasting antibodies by B cells, activate cytotoxic CD8^+^ T cells^3^, and directly eliminate virus-infected cells or tumor cells^4–6^. CD4^+^ T cells become active upon coming into contact with their corresponding antigens, which are presented on the surface of specialized antigen-presenting cells (APCs) via major histocompatibility complex (MHC) class II molecules^7^. The extensive activation and expansion of antigen-specific CD4^+^ T cells are critical for prolonged immune response and the establishment of immune memory^8^. Since CD4^+^ T cells are important components in inducing potent immune responses in various scenarios, including inflammation, viral infection, vaccination, cancer immunotherapy, and autoimmune diseases, the identification of peptide presentation by specific MHC II alleles is fundamental to understanding the underlying physiological biology of immune responses and for clinical applications.

Advances in delivery, stability, and design have shifted focus to peptide-based vaccines, enhancing immunogenicity and stability through epitope processing and post-translational modifications, with promising potential for chronic viral diseases and cancer treatment^9^. Due to the importance of decoding MHC II-presented epitopes, current methods include LC-MS/MS analysis of eluted peptides^10^ and computational prediction algorithms^11^. LC-MS/MS provides reliable data on naturally occurring peptides^12,13^, but lacks comprehensive coverage across diverse HLA types and has sensitivity limitations that may miss peptides with confirmed T cell reactivity^14^. While computational methods can identify strong MHC I binders^15^, they are less reliable for MHC II binders^16^. Recent improvements like NetMHCIIpan-4.1 show better prediction performance^17^. However, its suboptimal prediction performance might be due to the few lack of binding data for adequate training, especially for MHC II ligands. Taken together, a robust, high throughput assay to screen peptide-MHC II interactions is urgently desired. Peptide vaccines against infectious diseases and cancer are designed to augment the specific T cell immune responses^18–20^. The effectiveness of antibody-based protection is closely linked to the functionality of specific viral antigens, limiting the range of targets and increasing the likelihood of the pathogen evolving to escape immune responses^21,22^. Conversely, T cell-mediated protection relies primarily on the recognition of pathogen-derived peptides presented by MHC molecules, and it is relatively independent of the physiological function or location of the target protein. As a result, while only certain epitopes on surface proteins are susceptible to neutralizing antibodies, there is a much broader range of peptides that can serve as targets for T cells. This expanded epitope diversity provides a larger space for therapeutic development.

Here, we designed a rapid and high-throughput assay, namely, Exploring Landscape assay of Immunogenic T cell Epitopes for MHC II (EliteMHCII), to identify the landscape peptides from T cell epitopes across with global prevalent HLA alleles. Our approach allows robust and precise HLA-II prediction for immunodominant CD4^+^ T cell epitopes, generating a high-resolution landscape detailing peptides across various HLA alleles would be immensely beneficial to advance the research and therapeutic efforts.

Previous efforts to identify MHC II-restricted CD4⁺ T cell epitopes have led to the development of various platforms, including multimer-based staining systems^23^, yeast display technologies^24,25^, and mammalian epitope display frameworks^26^. However, each of these systems has limitations: multimer-based tools enable functional interrogation but generally lack quantitative resolution of peptide-MHC binding affinities; yeast display offers ultra-high throughput screening, yet cannot directly quantify binding or reliably reproduce human MHC II processing contexts; mammalian cell-based approaches enable peptide display but suffer from low scalability and assay complexity.

In contrast, we developed EliteMHCII as a quantitative, scalable, and mechanistically informed platform. This system couples biochemical affinity profiling across 24 globally prevalent HLA-DR alleles (covering 97.4% of the global population) with HLA-DM-mediated peptide exchange, enabling direct IC_50_ and K_i_ quantification for each peptide-HLA interaction (ranging from < 10 μM). Importantly, we further integrated tetramer-based functional readouts in vaccinated individuals and humanized mice, bridging biochemical binding strength with CD4⁺ T cell immunogenicity. These design principles are consistent with vaccine epitope selection criteria reported in a 2022 Nature study^19^, in which candidate peptides were prioritized based on the presence of detectable T cell responses in over 50% of SARS-CoV-2 convalescent individuals. Thus, EliteMHCII unites mechanistic binding resolution with functional immune validation in a single discovery-to-validation pipeline.

## Results

### Establishment of Exploring Landscape assay of Immunogenic T cell Epitopes for MHC II,EliteMHCII

To identify antigen-derived MHC II T cell peptide epitopes and quantify the relative affinity of the peptide epitopes with MHC II, we developed an Exploring Landscape assay of Immunogenic T cell Epitopes for MHC II (EliteMHCII). Firstly, to facilitate rapid and high-throughput allele MHCII protein production, knob into hole Fc tagged MHC II^27^ were expressed from Expi293F suspension culture **(Figure S1A)**. MHC II proteins were purified using Ni beads, followed by size-exclusion chromatography **(Figure S1B)**. Negatively stained electron microscopic analyses further verified their compositional integrity and conformational states **(Figure S1C)**. According to the allele frequency data sourced from the Allele Frequency Net Database (AFND)^28^, the most frequency of 24 common HLA-DR alleles across global populations were selected, representing a total of 83.9% of the global population, and covering 97.4% of individuals with at least one allotype **(Figure 1B)**. To demonstrate epitope loading, we developed a saturation binding assay to load antigen binding cleft with biotin-labeled human Class II-associated Invariant Chain Peptide (CLIP) **(Figure S1D)**. The saturation binding affinity (EC_50_) of human CLIP and 24 HLA-DR **(Figure 2A)**, were detected using our assay.

**Figure 1.**
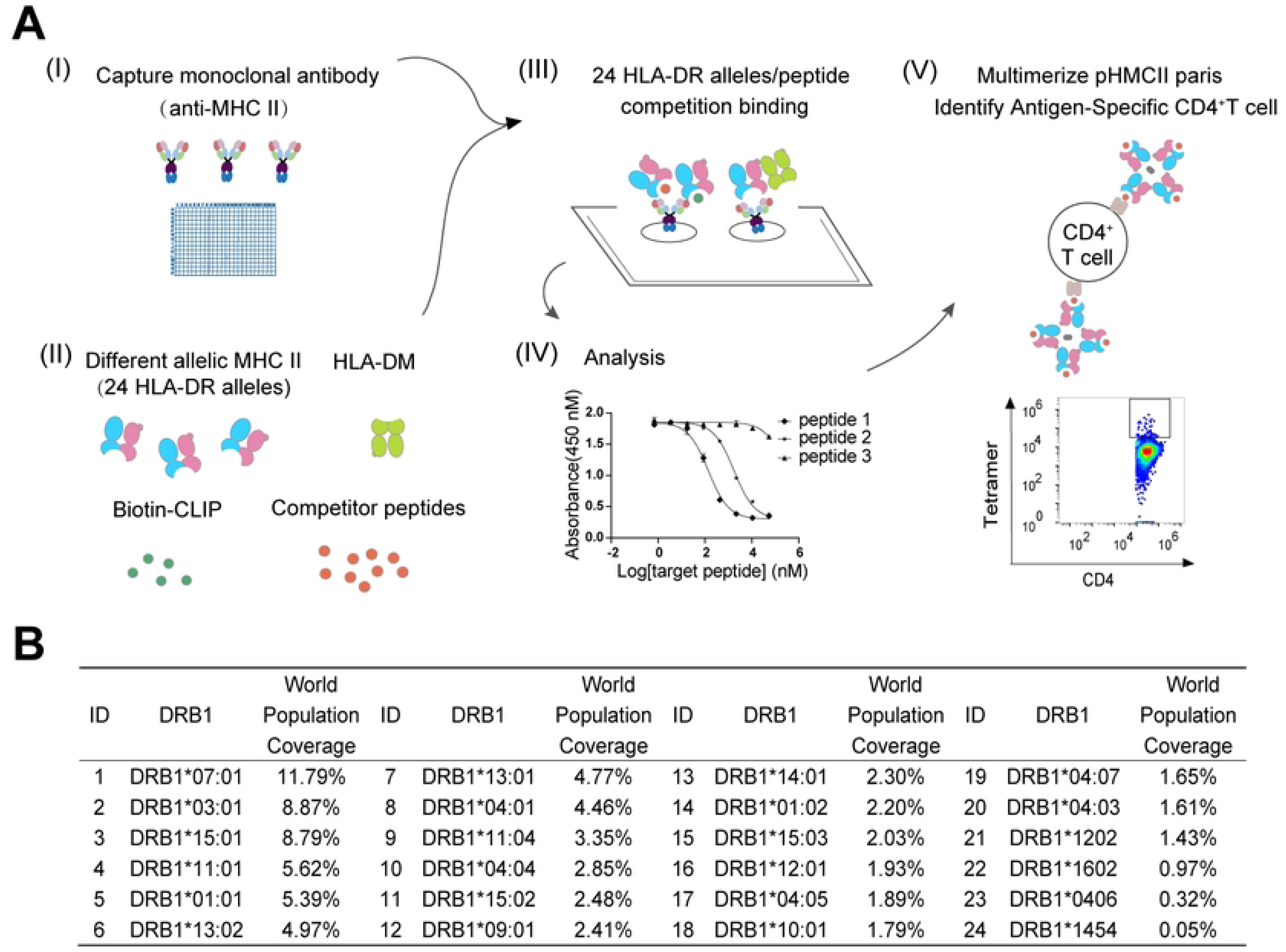
Establishment of an ex vivo rapid, efficient, stable, and quantitative system for MHC-II and peptide interactions. (A) Schematic overreview of EliteMHCII assay, the competitive binding of MHC-II molecules and antigenic peptides. The detection system includes catalysis, stabilization of MHC-II conformation, peptide editing by HLA-DM, bio-CLIP probe, and detection of target antigenic peptides, enabling rapid affinity detection for various MHC-II and antigenic peptides. (B) The coverage frequency of top 24 DRB1 alleles among world population, including DRB1*07:01, DRB1*03:01, DRB1*15:01, DRB1*11:01, DRB1*01:01, DRB1*13:02, DRB1*13:01, DRB1*04:01, DRB1*11:04, DRB1*04:04, DRB1*15:02, DRB1*09:0, DRB1*14:01, DRB1*01:02, DRB1*15:03, DRB1*12:01, DRB1*04:05, DRB1*10:01, DRB1*04:07, DRB1*04:03, DRB1*12:02, DRB1*16:02, DRB1*04:06, DRB1*14:54.

**Figure 2.**
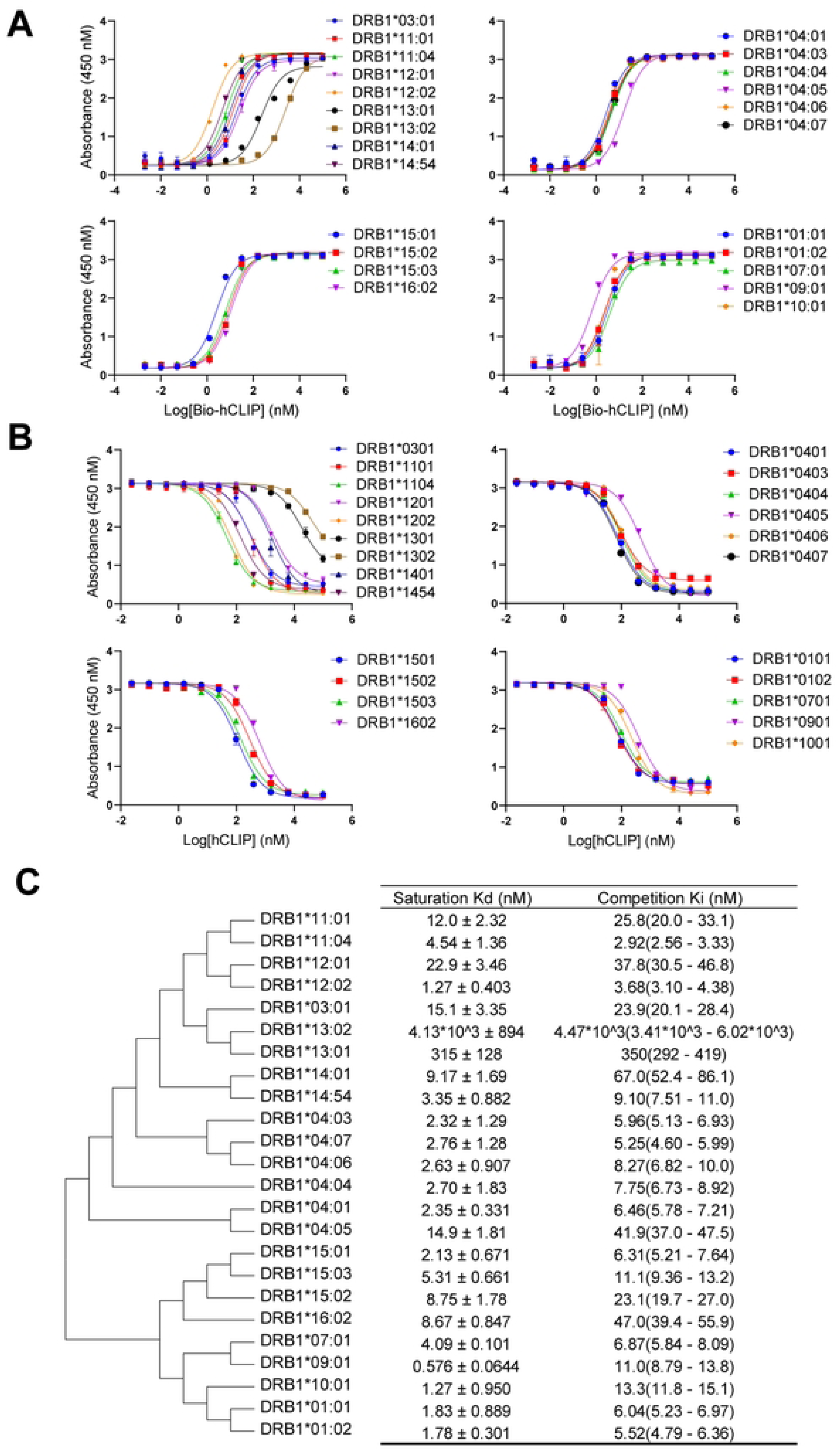
Binding of MHC II to the invariant chain-derived peptide, is regulated by allelic polymorphism in Class II. (A) The saturation binding affinity (EC_50_) of CLIP and 24 HLA-DR, including DRB1*01:01, DRB1*03:01, DRB1*04:01, DRB1*04:04, DRB1*04:05, DRB1*07:01, DRB1*09:01, DRB1*10:01, DRB1*11:01, DRB1*10:04, DRB1*12:01, DRB1*13:01, DRB1*15:01, DRB1*150:02, DRB1*15:03, DRB1*16:02, were detected using the EliteMHCII assay. Data representative of n = 3 independent experiments, each with n = 3 technical replicates. Data presented as mean A450 absorbance (circles) with standard deviation shown as error from n = 3 technical replicates. (B) Dose response IC_50_ curves of CLIP and 24 HLA-DR, including DRB1*01:01, DRB1*03:01, DRB1*04:01, DRB1*04:04, DRB1*04:05, DRB1*07:01, DRB1*09:01, DRB1*10:01, DRB1*11:01, DRB1*10:04, DRB1*12:01, DRB1*13:01, DRB1*15:01, DRB1*150:02, DRB1*15:03, DRB1*16:02, were detected using the EliteMHCII assay. Data representative of n = 3 independent experiments, each with n = 3 technical replicates. Data presented as mean A450 absorbance (circles) with standard deviation shown as error from n = 3 technical replicates. (C) Evolutionary tree analysis of 24 HLA-DR. Data presented as mean saturation binding affinity and competition binding affinity with standard deviation shown as error from n = 3 biological repeat. The Ki values obtained from competitive binding of CLIP and 24 HLA-DR.

To mimic the peptide exchange process that occurs within major histocompatibility complex class II-containing (MIIC) compartments, a competitive binding between MHC II and biotinylated human CLIP as probes was used. The dissociation of CLIP from MHC II was facilitated by HLA-DM, which accelerated the peptide exchange^29^ **(Figure 1A)**. The K_i_ values from the competitive binding (IC_50_) of CLIP and 24 HLA-DR **(Figure 2B)**, are comparable to the K_d_ values of saturation binding and those previously reported ^30–32^ **(Figure 2C)**.

Using this detection system, all 24 expressed HLA-DR variants were shown to bind with human CLIP, consistent with its crucial role^29,33^ in regulating the folding, trafficking, and peptide loading of major histocompatibility complex class II molecules. The evolutionary analysis of these 24 HLA-DR allotypes can be categorized into four major groups **(Figure 2C)**. The further apart the MHC II molecules are on the evolutionary tree, the greater the difference in the binding sites of CLIP. The majority of HLA-DR affinities fall within the 0-25 nM range. DRB1*13:01 and DRB1*13:02 exhibit relatively low affinities (below 300 nM), with the affinity of DRB1*13:01 for CLIP being approximately 10 times higher than that of DRB1*13:02. The difference in binding affinities between various MHC II alleles and CLIP indicates that the binding of MHC II to CLIP is regulated by allelic polymorphism within MHC II.

These results confirmed the robust binding of all 24 tested HLA-DR molecules to CLIP, validating the platform’s applicability for high-throughput peptide screening across globally representative alleles.

Compared to existing approaches^23–26^, EliteMHCII provides distinct advantages in quantifiability, allele breadth, and functional integration. Yeast display systems allow large-scale peptide screening but typically yield relative enrichment scores rather than absolute affinity metrics. Mammalian display approaches assess peptide presentability but depend on surrogate expression markers and lack K_i_ or IC_50_ data. Multimer-based staining enables direct T cell detection but omits upstream affinity screening, making it difficult to distinguish strong from weak epitopes before testing.

In contrast, EliteMHCII employs both saturation and competition assays to generate accurate binding curves for each peptide-HLA pair in a soluble protein context. This dual capability facilitates rigorous affinity comparison across alleles and allows rational prioritization of candidates for further immunological evaluation, providing a direct pipeline from biophysics to immune relevance.

### Molecular Dynamics Simulation of Affinity Differences Between HLA-DR and CLIP

Remarkably, compared to the majority MHC II, with the affinity within the 0-25 nM range, DRB1*13:01 and DRB1*13:02 exhibit relatively low affinities (300 and 4000 nM). To explain the differences in affinity observed in the platform assays, molecular dynamics (MD) simulations were employed (**Figure 3A-B**). The binding free energy (ΔG_bind_) was −39.94 kcal/mol for DRB1*13:01 and −38.24 kcal/mol for DRB1*13:02 (**Figure 3C**), indicating marginally stronger binding to DRB1*13:01. Analysis of the interaction energy components revealed that electrostatic energy (ΔE_ele_) played a dominant role in the binding process, with a significantly larger contribution compared to van der Waals energy (ΔE_vdW_) shown in **Table S1**.

**Figure 3.**
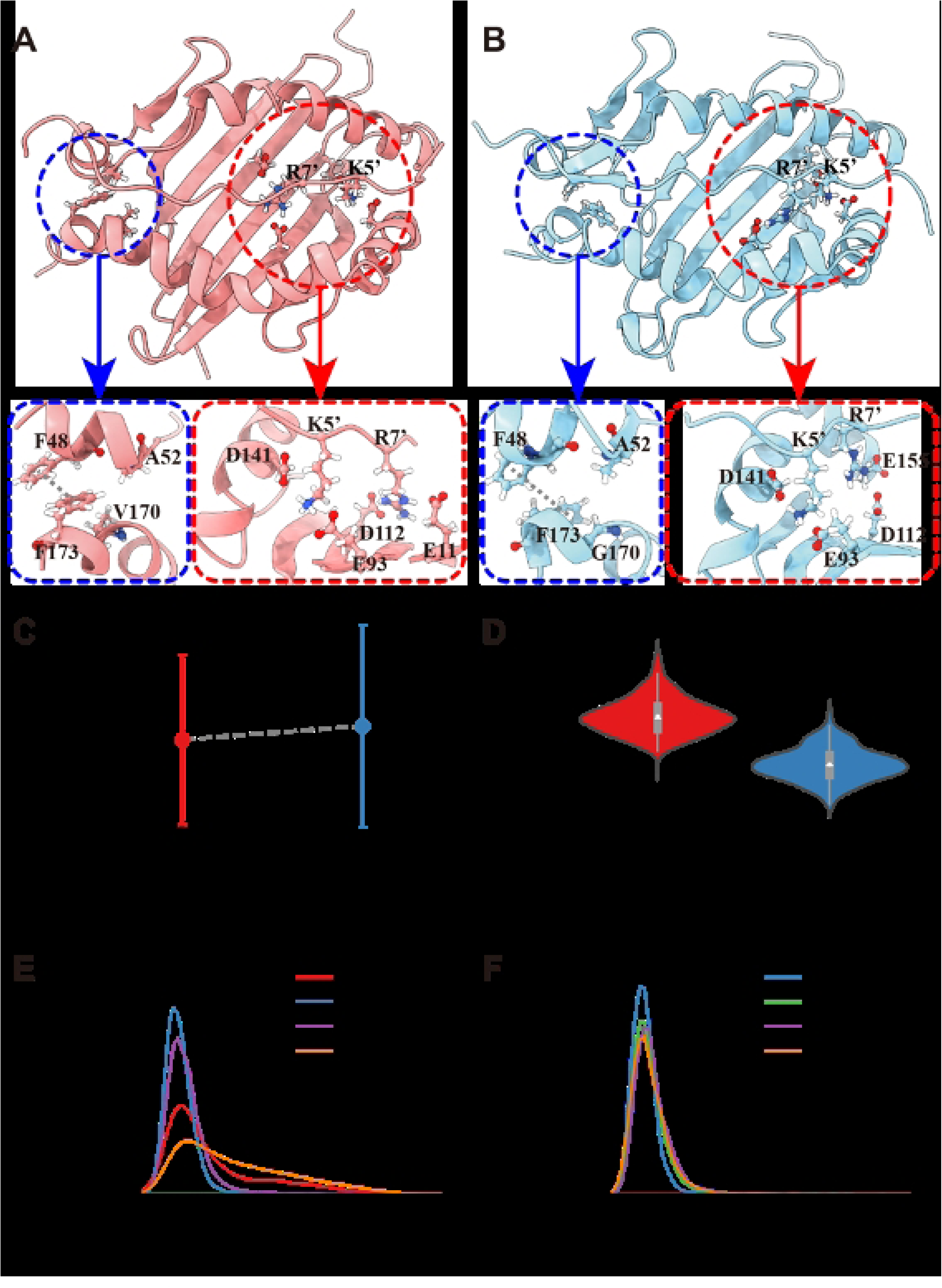
Molecular dynamics simulation analysis of CLIP peptide binding to DRB1*13:01 and DRB1*13:02 proteins at pH 7.4. Representative snapshot of CLIP peptide bound to (A) DRB1*13:01 and (B) DRB1*13:02 proteins. (C) Binding free energy (ΔGbind) of CLIP peptide with DRB1*13:01 and DRB1*13:02 proteins. Error bars represent standard deviation. (D) Violin plots showing the distribution of Voronoi volumes of hydrophobic core region in DRB1*13:01 and DRB1*13:02 proteins. The white dots represent the average volume, and the box plots show the interquartile range. The distribution of distances between the nearest heavy atoms of sidechains for different residue pairs in (E) DRB1*13:01 and (F) DRB1*13:02. R7’ and K5’ denote residues of the CLIP peptide.

The structural analysis revealed distinct differences between the two proteins. The hydrophobic core, formed by residues F48, A52, V170/G170, and F173 residues, exhibited a significantly reduced volume in DRB1*13:02 compared to DRB1*13:01 (**Figure 3D**). This reduction is primarily attributed to the V170G mutation in DRB1*13:02 (**Figure S2B**), resulting in a less tightly packed hydrophobic core. The mutation caused the F173 residue sidechain to move inward, disrupting the packing between F173 and F48 and causing the F48 sidechain to shift outward in DRB1*13:02 (**Figure S3B & S3D**).

The binding trend of the CLIP peptide with both HLA-DR proteins is similar, but the amino acid arrangement shows misalignment and twist. The binding orientation of the CLIP peptide also differed between the two proteins. In DRB1*13:01, the sidechain of Q15 residue of peptide pointed away from the F173 residue, whereas in DRB1*13:02, it pointed towards F173 residue due to the reduced hydrophobic core volume (**Figure S4**). This orientational difference resulted in the R7 residue of peptide (R7’) formed hydrogen bonds with D112 and E11 residues in DRB1*13:01, while it formed hydrogen bonds with D112 and E155 residues in DRB1*13:02 (**Figure 3E-F**). The K5 residue of the peptide (K5’) primarily interacted with E93 residue in both proteins, with an additional interaction with the D141 residue observed in DRB1*13:02. A strong hydrogen bond pair between R76 and D141 residues formed in both DRB1 proteins, establishing a robust gating that restricted the binding regions of K5’ residue and E93 residue (**Figure S5F**). These structural differences resulting from the V170G mutation, contribute to the slightly stronger binding affinity of the CLIP peptide to DRB1*13:01 compared to DRB1*13:02. The MD simulation results are consistent with experimental observations.

### EliteMHCII for Peptide Screening

To evaluate the feasibility of the platform for screening peptides with high affinity for a specific protein, the binding affinities of 53 peptides from the receptor binding domain (RBD) of SARS-CoV-2 were screened as an example. The 53 peptides were synthesized from the SARS-CoV-2 RBD (amino acids 319–541) as 15-mer peptides with a 4-amino acid overlap. The RBD is the region of the spike protein that binds to the ACE2 receptor for viral entry.

Previous studies^22^ have categorized peptides with a binding affinity greater than 10 µM as either binding or non-binding. Here we defined the affinity of less than 100 nM as high affinity, between 100 nM and 1 μM as moderate affinity, and between 1 μM and 10 μM as low affinity. In our study, the lowest proportion of RBD peptides with a binding affinity above 10 µM for HLA-DR was observed for DRB1*0102, at under 10%, followed by DRB1*04:03, DRB1*13:02, DRB1*15:03, and DRB1*12:02, with all these alleles showing proportions below 30%. This suggests that these HLA-DR alleles have lower affinity for most RBD peptides, indicating weaker peptide presentation capabilities. On the other hand, DRB1*07:01, DRB1*01:01, DRB1*04:05, DRB1*10:01, and DRB1*16:02 showed a proportion of RBD peptides with a binding affinity above 10 µM exceeding 60%, suggesting that these alleles might have superior capability to present RBD peptides. The heat map suggesting some HLA subtype might have a poor ability or have superior capability to present RBD peptides.

To identify the potential peptides with high immunogenicity across different populations, we aimed to select the candidate peptides that exhibited both high binding affinity and broad compatibility with the most popular HLA-DR alleles within the RBD. The HLA-DR affinity score for each RBD peptide across 24 HLA-DR alleles was determined by combining the frequency of different HLA-DR groups. Our data showed that RBD_323-337_, RBD_339-353_, RBD_447-465_, and RBD_511-525_ peptides were capable of binding HLA-DRs with a 50% coverage of the population in a high or moderate affinity manner. Interestingly, high or moderate binding affinity peptides with more than 20% of global HLA-DR haplotypes were clustered together within RBD region, including RBD_323-337_, RBD_339-361_, RBD_363-389_, RBD_427-445_, RBD_443-465_, RBD_483-505_, and RBD_499-529_, with a significant portion demonstrating high affinity binding to over 10% of the population. Conversely, there were low-affinity epitope clusters including RBD_411-433_, RBD_435-453_, and RBD_475-493_. Notably, the epitope RBD_419-433_ fails to bind to any of the 24 tested HLA-DR types. Interestingly, the above peptides with broad binding to HLA-DR with high affinity were clustered distributed across the RBD region **(Figure 4B)**. Therefore, our platform can identify regions within a protein that exhibit high affinity for HLA-II alleles.

**Figure 4.**
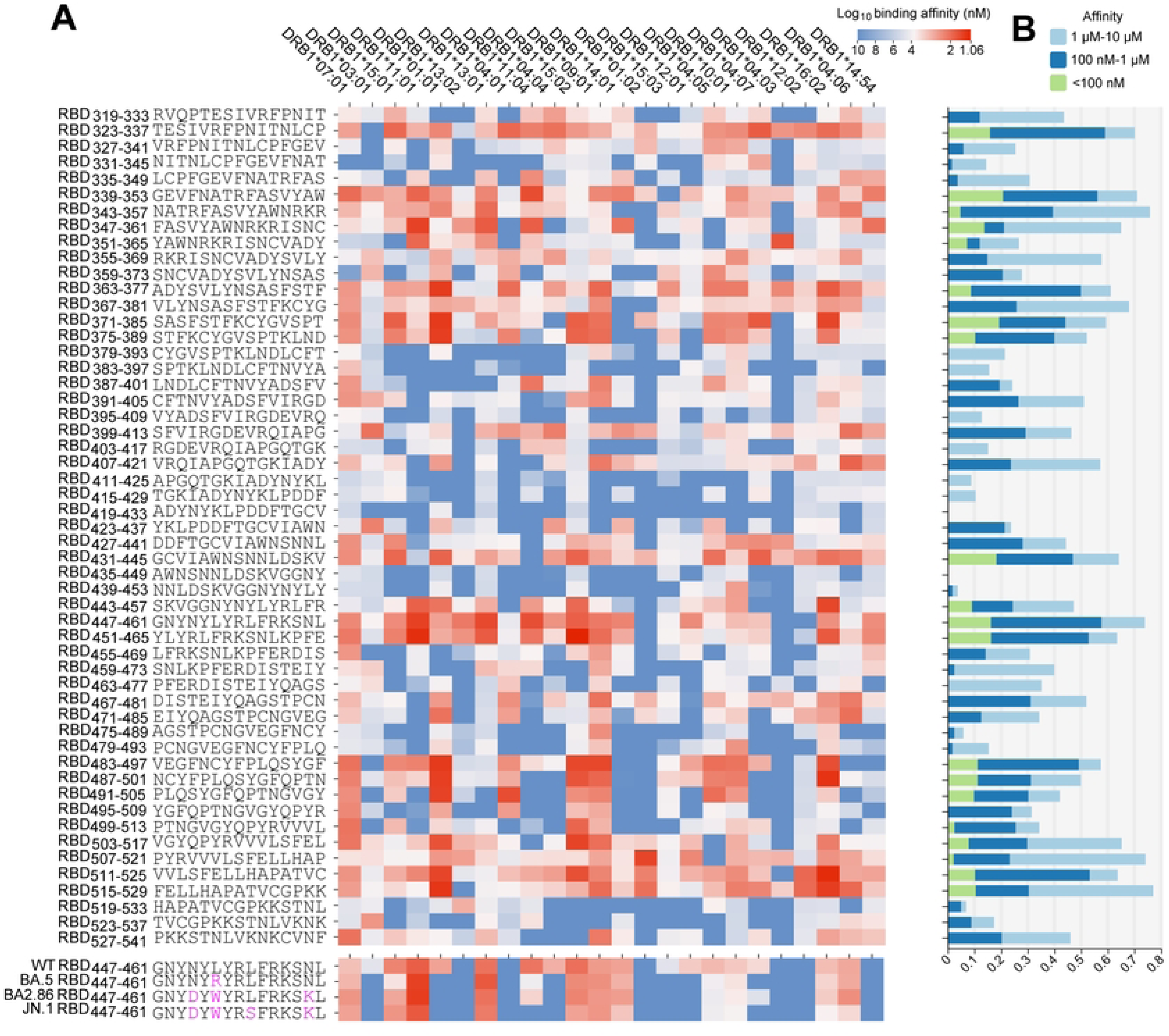
Validation of EliteMHCII assay using SARS-CoV-2 RBD derived HLA-DR-binding peptides. (A) A heatmap is shown summarizing the distribution of binding affinity between HLA-DRs and RBD peptides. The color represents the degree of binding affinity, in which high binding affinity was shown in dark read, whereas the weak affinity was shown in dark blue. The Heatmap also showing the binding affinity of RBD_447-461_ peptide from SARS-CoV-2 variants including wildtype strain, BA.5, BA.2.86, and JN.1 with HLA-DR alleles. The mutated aminio acids were highlighted in purple. (B) Affinity score of 53 overlapping RBD peptides with high-frequency HLA-DR alleles based on their binding affinity as well as HLA allele frequencies. The affinity score was calculated

To assess whether the emerging variants might lead to immune escape for CD4^+^ T cell responses, we determined the binding affinity of emerging variants’ specific epitope with HLA-DR. We focused on the RBD_447-461_ epitope and the sequences at the RBD_447-461_ were analyzed for the wide type (WT) strain, Omicron BA.5, BA.2.86, and JN.1 variants. This epitope was selected to study the impact of the L455S recent mutation in the JN.1 variant on epitope binding. **(Figure 4A)**. As the EliteMHCII assay indicated, the binding affinity of 11 or 12 HLA-DRs and RBD_447-461_ derived from BA.5, BA.2.86, and JN.1 variants were significantly reduced, compared to RBD ^WT^. RBD ^JN.1^ exhibited the lowest affinity with 12 HLA-DRs, compared to RBD ^BA.5^and RBD ^BA.2.86^. The epitopes RBD of BA.4/BA.5, BA.2.86, and JN.1 variants lost their binding affinity to DRB1*03:01, DRB1*01:01, DRB1*04:04, DRB1*12:01, and DRB1*04:07, compared to RBD ^WT^. In the case of JN.1, RBD_447-461_ epitope completely lost its binding capability to DRB1*15:01 and DRB1*11:04. Overall, RBD_447-461_ from Omicron variants might result in the loss of presentation by these HLA-DR molecules. Moreover, the binding affinity of RBD ^BA.5^ and RBD ^BA.2.86^ to DRB1*11:04 and DRB1*04:05 has significantly declined by over 16-fold, compared to RBD ^WT^. In particular, 6 HLA-DRs, including DRB1*04:01, DRB1*01:02, and DRB1*15:03, showed no binding for either WT strain or the emerging variants, while the other 6 HLA-DRs maintained binding capability with the WT strain and the tested Omicron variants **(Figure 4A)**. Reduced binding affinity of epitopes to HLA-DR molecules impairs their presentation to CD4^+^ T cells, potentially leading to immune evasion. The significant decrease in binding affinity, particularly in variants like JN.1, may prevent effective immune recognition and response. Therefore, this assay can help identify key mutations within CD4^+^ T cell epitopes that may potentially impact virus immune escape.

### Comparison of EliteMHCII and NetMHCIIpan

To evaluate the relationship between the predicted binding affinities from NetMHCIIpan-4.1 and the experimentally measured values from our EliteMHCII assay, we performed correlation analysis across various HLA-DR alleles. The results showed the binding affinities predicted by NetMHCIIpan-4.1 algorithm partially correlated with that from our EliteMHCII assay, with the correlation co-efficiencies ranging from 0 to 0.7 **(Figure 5A)**. Specifically, for 16 DRB1 alleles including DRB1*01:01, DRB1*07:01, and DRB1*11:01, their binding affinity were highly correlated with the predicted affinity (r≥0.5, p≤0.0006). For DRB1*14:54, DRB1*14:01, DRB1*04:07, and DRB1*04:06, they have moderate correlation (0.3≤r < 0.5, p≤0.05). Nevertheless, the binding affinity of DRB1*12:02, DRB1*0403, DRB1*1302, and DRB1*0102, were not correlated with the predicted affinity (p > 0.05) **(Figure 5B)**. Our results indicated the promiscuity of HLA-DR alleles in recognizing peptides across diverse motifs, which further highlight the importance of our platform to generate abundant and high-fidelity training datasets.

**Figure 5.**
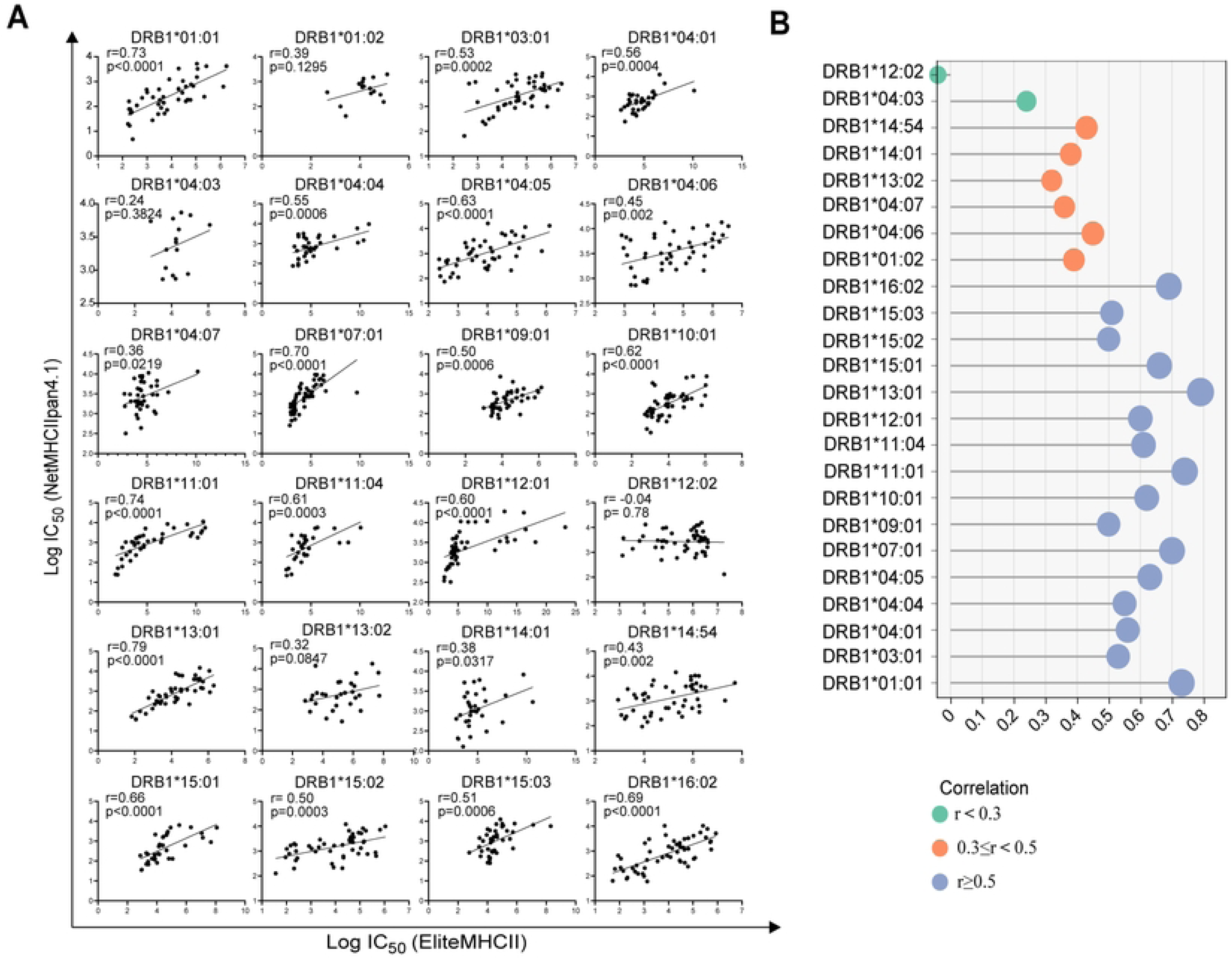
Benchmarking EliteMHCII performance against existing Mass spectrometry analysis and computational prediction algorithm. (A) and (B) Correlation analysis of log10-transformed binding affinity (IC_50_) between RBD peptides and 24 HLA-DR detection and predicted affinity by NetMHCIIpan-4.1. 16 DRB1 alleles, including DRB1*01:01, DRB1*03:01, DRB1*04:01, DRB1*04:04, DRB1*04:05, DRB1*07:01, DRB1*09:01, DRB1*10:01, DRB1*11:01, DRB1*10:04, DRB1*12:01, DRB1*13:01, DRB1*15:01, DRB1*150:02, DRB1*15:03, DRB1*16:02, showed a high correlation (r≥0.5, p≤0.0006) between their binding affinity and the predicted affinity. The binding affinity of DRB1*14:54, DRB1*14:01, DRB1*04:07, DRB1*04:06 measured by EliteMHCII assay was moderately correlated with NetMHCIIpan-4.1 (0.3≤ r<0.5, p≤0.05). The binding affinity of DRB1*12:02, DRB1*04:03, DRB1*13:02, and DRB1*01:02 was not correlated with the predicted affinity (p > 0.05).

### EliteMHCII demonstrates proficient presentation of immunogenic CD4^+^ T cell epitopes

To validate the immunogenicity of epitope peptides screened by the EliteMHCII assay, 10 peptides (RBD_323-337_, RBD_371-385_, RBD_399-413_, RBD_419-433_, RBD_423-437_, RBD_431-445_, RBD_447-461_, RBD_483-497_, RBD_511-525_, and RBD_523-537_) were used to identify and characterize SARS-CoV-2 RBD specific CD4^+^ T cell responses from vaccinated volunteer. The selection of these 10 peptides was aimed at ensuring the inclusion of a diverse range of HLA-DR alleles, covering peptides with high, medium, low binding affinities, as well as non-binding peptides. Each HLA-DR allele corresponds to both binding and non-binding peptides. Some peptides exhibit a balanced distribution of binding and non-binding across 9 HLA-DR alleles, while others are biased towards either predominantly binding or predominantly non-binding **(Figure S7)**. This approach enables a more comprehensive assessment of the immunogenicity of peptides associated with different HLA-DR alleles. And antigen-specific CD4^+^ T cells from peripheral blood mononuclear cells (PBMCs) of the eight volunteers (HLA-DR allotypes are shown in Table S5) were identify by the fluorescently labeled peptide-HLA-DR tetramers.

To find out whether there are CD4^+^ T cells specific to high-affinity and low-affinity peptides in volunteers who have been immunized with the vaccine, the affinity of 10 peptides to DRB1*07:01 and DRB1*12:01 of donor 1 was compared with the proportion of epitope-specific CD4^+^ T cells detected in PBMCs. Consistent with our previous result that three peptides with high affinity for DRB1*07:01, including RBD_323-337_, RBD_483-497_, and RBD_511-525_, we also detected a positive population of CD4^+^ T cells which is specific to the above peptides, whereas we did not detect the positive CD4^+^ T cell responses specific to the remaining three peptides. Additionally, all ten peptides with high or low affinity for DRB1*12:01 corresponded to the proportion of peptide-specific CD4^+^ T cells detected by tetramers **(Figure 6A, Figure S8A)**. Thus, high-affinity peptides identified by the EliteMHCII assay correlated with CD4^+^ T cell responses, while low-affinity peptides showed limited or no T cell responses, highlighting the assay’s ability to predict immunogenic peptides.

**Figure 6.**
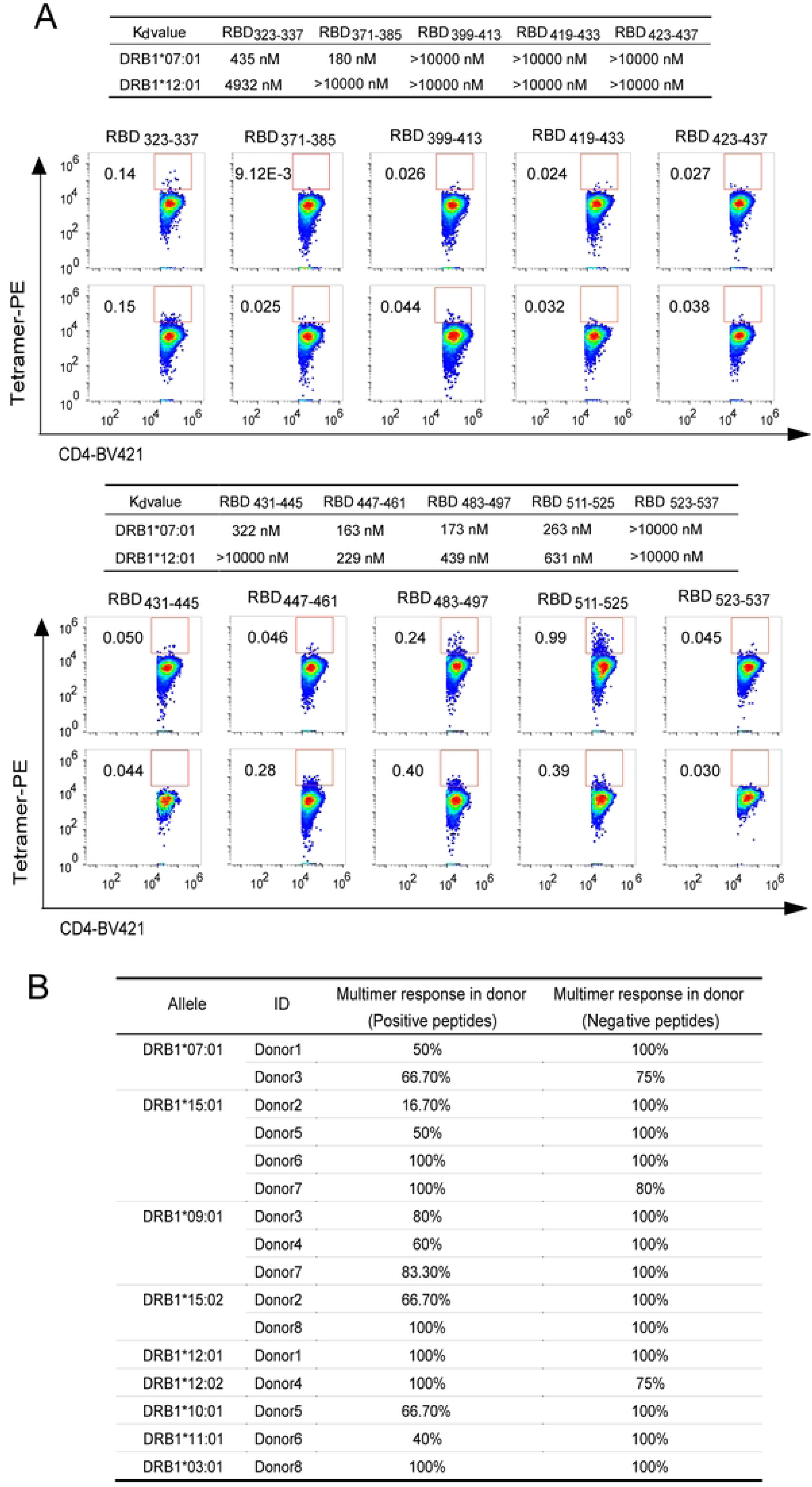
The flow cytometry analysis of HLA-DR tetramer staining for peptide-specific CD4^+^ T cell responses. (A) Binding Affinity (K_d_) of 10 indicated RBD peptides with with DRB1*07:01 and DRB1*12:01, as measured by EliteMHCII assay, including RBD_323-337_, RBD_371-385_, RBD_399-413_, RBD_419-433_, RBD_423-437_, RBD_431-445_, RBD_447-461_, RBD_483-497_, RBD_511-525_, RBD_523-537_. Flow cytometry analysis for RBD peptide-specific CD4^+^ T cells from PBMC collected two weeks post vaccination using DRB1*07:01 and DRB1*12:01 tetramer staining. (B) Detection rates of specific CD4^+^ T cells using tetramers for each positive peptide and negative peptide (a total of 10 peptides) across 10 different HLA-DR types from 8 donors.

To analyze the correlation between the affinity and immunogenicity of the peptides. we statistically analyzed the affinity and detected proportion of CD4^+^ T cells for the 10 peptides across the eight volunteers **(Table S3)** with nine different HLA-DR allotypes. The concordance rate for low-affinity peptides was greater than 75%, and for high-affinity peptides, there was individual variability with a concordance rate higher than 66% in 11/16 **(Figure 6B)**. Statistical analysis demonstrated a high concordance rate between peptide binding affinity and CD4^+^ T cell responses **(Figure 6B)**.

Overall, the binding affinity identified by EliteMHCII assay was highly correlated with the immunogenicity of these peptides, as indicated by CD4^+^ T cell responses. This indicates our EliteMHCII assay effectively identifying immunogenic high-affinity MHC-II binding peptides across diverse HLA-DR allotypes.

### Evaluation of peptide vaccine immunogenicity in humanized mouse models

The binding affinity of pathogen-derived peptides to MHC II dictates their antigenicity, peptides with higher affinity to MHC II are potentially prone to trigger the T cell immune response^33^. Previous studies reported that SARS-CoV-2 CD4^+^ T cell response positively correlates with higher antibody titer and persistence of neutralizing antibodies in patients^34,35^. To interrogate whether high-affinity peptides can induce a more potent CD4^+^ T cell response and benefit the production of antibodies, humanized immune system mice reconstituted by hematopoietic stem and progenitor cells (HSPCs) from a donor with HLA-DRB1*01:01 and HLA-DRB1*14:54 haplotype were immunized with high-affinity peptides (RBD_399-413_, RBD_431-445_, RBD_447-461_) and low-affinity peptides (RBD_395-409_, RBD_475-489_, RBD_499-513_), respectively. And before immunization, hydrodynamic injection of GM-CSF and IL4 plasmid were performed to stimulate the development of dendritic cells (DCs) and promote the T and B cell response^36^. One week after primary peptide vaccination, we administered mice with 4RBD-Fc that can elicit a strong B-cell immune response in K18-hACE2 mice (unpublish data). A peptide vaccine boosters with the same formula were given to mice on day 14 **(Figure 7A)**. 6 days after the last dose vaccination, we detected significantly higher serum antibody titer in mice immunized with high-affinity peptide vaccination compared with the average antibody titer of mice receiving low-affinity peptide vaccination **(Figure 7B)**. We also isolated the splenocytes from humanized mice and identified the peptide-specific CD4^+^ T cells with HLA-DR tetramers loaded with high-affinity and low-affinity peptides **(Figure 7A and Figure S8B).** Indeed, we observed the high frequencies of peptide-specific CD4^+^ T cells against RBD_431-445_ and RBD_447-461_ in the high-affinity peptide immunized mice **(Figure 7C)**. In contrast, the presence of peptide-specific CD4^+^ T cells for the low-affinity peptides was nearly undetectable **(Figure 7C)**. Overall, the mice receiving high-affinity peptides showed a tendency with higher CD4^+^ T cell response. Concurrently, the high frequency of activated peptide-specific CD4^+^ T cells was noted after *ex vivo* stimulation with corresponding peptides, as indicated by the T cell activation induced marker (AIM) CD69 and CD25^37,38^ **(Figure 7D**. Taken together, the high-affinity peptides revealed by our EliteMHCII assay have a high potential to trigger stronger immune responses in the humanized mice model.

**Figure 7.**
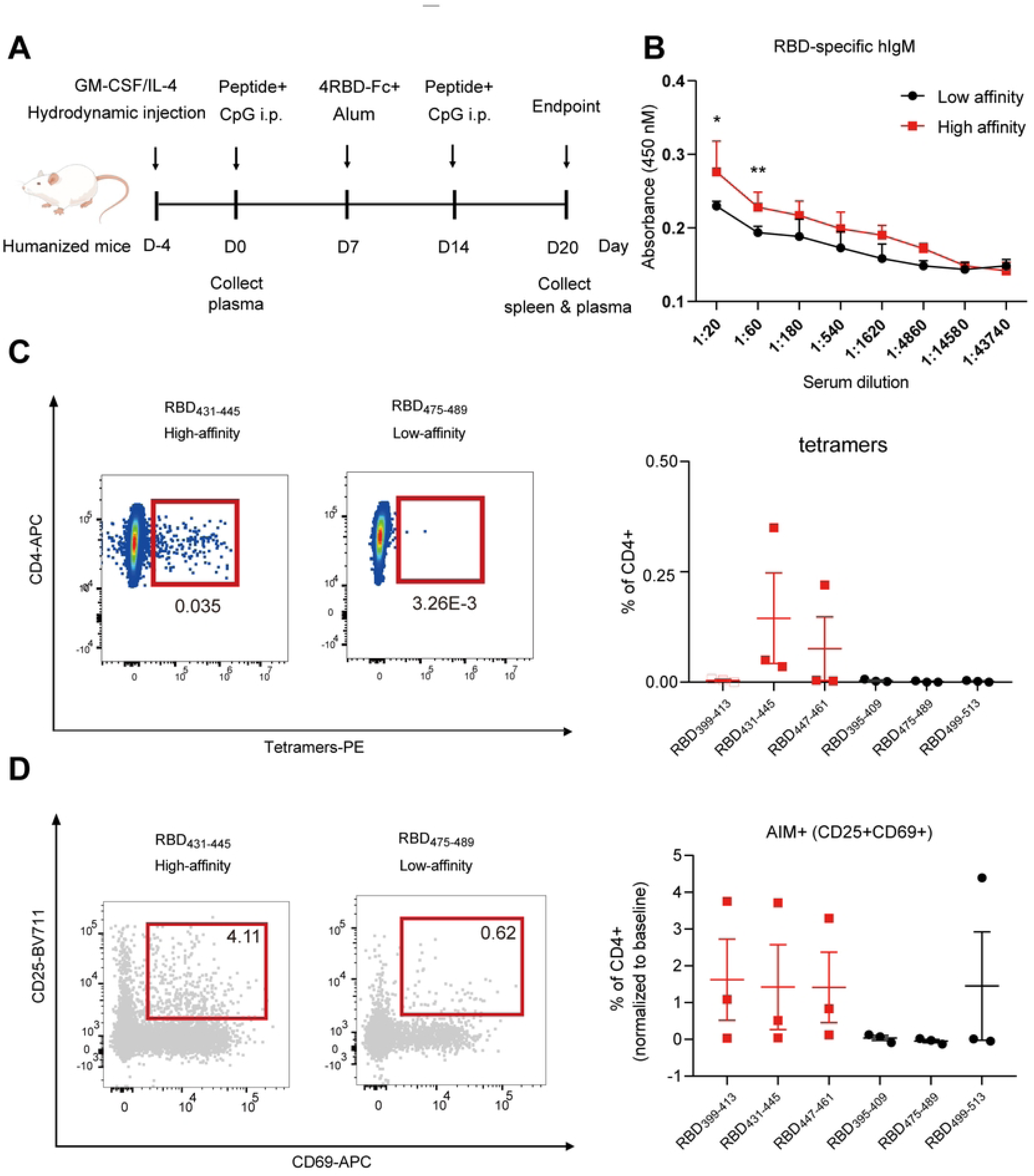
RBD epitope specific T-cell responses induced by high-affinity and low-affinity SARS-CoV-2 RBD peptides in humanized mice model. (A) Immunization strategy of peptide vaccines for humanized mice. Humanized mice were immunized with the peptide vaccine containing high-affinity peptides (RBD_399-413_, RBD_431-445_, and RBD_447-461_) and low-affinity peptide candidates (RBD_395-409_, RBD_475-489_, and RBD_499-513_), respectively. All mice were administered 4RBD-Fc plus alum adjuvant on day 7. Subsequently, mice were given a booster with the same formula on day 14. The mice spleen and serum plasma were collected on day 20. (B) RBD-specific IgM antibody titer from the groups receiving high-affinity and low-affinity peptides, respectively. (C) Representative flow cytometry plot for RBD epitope tetramer-specific CD4^+^ T cells in mice immunized with either high-affinity peptides or low-affinity peptides 20 days post-immunization (Left). Bar plot summarized RBD^+^CD4^+^ T cells against the indicated peptide, and each dot indicated an individual mouse (Right). (D) Representative flow cytometry plot for CD69^+^ CD25^+^ population in CD4^+^ T cells after *ex vivo* stimulation with cognate peptide (Left). Bar plot summarized the frequency of CD69^+^ CD25^+^ in CD4^+^ T cells against the indicated peptide, and each dot indicated an individual mouse (Right)

## Discussion

### EliteMHCII offers improvements in efficiency, sensitivity, throughput, and feasibility

Presentation of antigenic peptides by MHC-II proteins determines T helper cell reactivity. Accurate prediction of antigen presentation by human HLA class II molecules would be valuable for vaccine development and cancer immunotherapies. Nevertheless, a comprehensive understanding of MHC II epitopes is lacking due to their complexity compared to MHC I ligands^39^. Given the difficulty in accurately predicting important and immunodominant HLA-II epitopes through *in silico* assessment along with peptide-HLA-II binding, here we developed a robust, high-throughput platform for determining the binding affinity of the MHC class II and the antigen of our interest.

In our established EliteMHCII assay, we tackled several challenges in this field. Firstly, EliteMHCII assay was conducted in a high throughput manner, which could automatically detect 384 samples at one time. Therefore, our platform allows a rapid identification of the immunodominant T cell epitopes among hundreds of peptides within 48 hours, compared to 1 month using traditional mass spectrometry approaches, which could greatly accelerate the development of peptide-based vaccines in the setting of emerging infectious diseases and personalized cancer immunotherapy. Secondly, we expressed 24 HLA-DR proteins which covered 83.92% of the global population. Although emerging approaches such as yeast display have been developed to quickly identify the MHC-II restricted T cell epitopes, the number of studied MHC-II alleles is very limited^24,25,40^. Thirdly, to display MHC-II molecules in their native, conformational heterodimeric form, we used mammalian cell Expi293F to express the soluble MHC-II proteins which were further validated by the conformational specific antibody L243. Fourthly, EliteMHCII assay achieved high sensitivity, which could detect the binding affinity as low as the nanomolar level. We observed that EliteMHCII generated significantly larger datasets compared to eluted ligand MS, suggesting a potentially enhanced performance in predicting peptide affinity. Therefore, our EliteMHCII assay extends and improves upon the current platforms that were limited by efficiency, sensitivity, throughput, feasibility, and scalability beyond the model alleles and epitopes.

Given the importance of understanding the impact of virus mutation on T cell responses, our EliteMHCII platform also provides a reliable tool to monitor the impact of emerging viral variants on the immune recognition of CD4^+^ T cells. Using our EliteMHCII assay, we showed that viral variants of HLA-II presented epitopes escaped the pre-existing immune memory. Our study highlights the consequences that viral evolution could have generally on the peptide epitopes presented to CD4^+^ T cells on the HLA-II platform. Additionally, since our EliteMHCII assay covers the most popular MHC II alleles, we could monitor the population immunity status by measuring the dynamics of the binding affinity of emerging variants with diverse MHC II alleles.

### Immunogenic epitope identification

Our EliteMHCII assay allows high-dimensional identification of immunogenic epitopes to prioritize vaccine candidates in diverse clinical scenarios, including infectious diseases, cancers, and autoimmune diseases. To validate our findings *in vivo*, we identified the potential immunogenic epitopes across most of the global population in the context of COVID-19. Here using SARS-CoV-2 RBD protein as an example, we found that the most of epitope immunogenicity correlated with peptide affinity identified from our EliteMHCII assay. Interestingly, a recent study reported that despite the high immunogenicity of S_486–505_ and S_511–530,_ both epitopes had weak binding affinity with DRB1*01:01 with IC_50_ of 3046 nM and 2624 nM, respectively^22^. Nevertheless, our assay showed that S_486–505_ (referring to RBD_487-501_) and S_511–530_ (referring to RBD_491-505_) had high binding affinity to DRB1*01:01 with IC_50_ of 293 nM and 152 nM, respectively. We speculate the disparity of binding affinity from two studies might be attributed to mammalian expression versus the *E.coli* expression system for the MHC II protein. It is possible that MHC II protein produced from the mammalian system is appropriately folded, which might represent the antigen presentation *in vivo*. Nevertheless, we also notice that there was a slight inconsistency between our assay and the immunogenicity of the peptides, which might be due to the fact that presentation of antigens is essential but not sufficient for the induction of robust T cell responses.

Notably, several high-affinity peptides induced strong CD4⁺ T cell staining signals in both vaccinated humans and humanized mice, while peptides with low binding affinity (K_i_ > 10 μM) failed to elicit detectable responses. Among the top-ranked binders, >50% triggered functional CD4⁺ T cell activation, whereas >75% of low-affinity peptides did not, consistent with the immunogenicity correlation seen in natural infection cohorts. These results were obtained at a time point approximately six months post-vaccination, when circulating antigen-specific T cells are expected to contract, possibly underestimating true response rates.

Unlike multimer-based platforms^23^ that lack affinity-based peptide stratification, or yeast^24,25^ and mammalian display^26^ systems that often require secondary functional assays, EliteMHCII integrates biochemical affinity profiling with direct T cell validation in a closed-loop format. This allows efficient and quantitative prioritization of immunogenic epitopes.

### Clinical application potential

EliteMHCII also has a great potential for clinical application in cancer immunotherapy. Recent neoantigen-based cancer immunotherapy requires identification of tumor derived mutation, prediction of peptide binding with HLA molecules, peptide synthesis, and immunogenic neoantigen screening, which takes a median of 100 days as indicated by recent clinical trials. In contrast, a library could be constructed using our EliteMHCII platform, which includes the tumor-specific mutated peptides specific to diverse MHC II alleles. Given the two HLA-DR alleles from an individual patient, EliteMHCII could quantify the binding affinity of hundreds of peptides with within 3 days. Therefore, our platform provides a platform for timely and efficient discovery of individually tailored neoantigen peptide, potentially providing a feasible strategy for developing precision immunotherapeutic strategies.

Peptide-based vaccine has its intrinsic limitations, including limited HLA restriction, rapid epitope identification, and broad coverage for global popular MHC II alleles, while our EliteMHCII overcomes these limitations by functioning as a high throughput platform to rapidly identify the human HLA-II immunopeptidome or utilizing this platform for immunogenicity prediction.

### Limitations

Our study has several limitations. First, we set out during the pandemic to use the HLA-DR1 allotype as a model to assess the presentation and recognition of SARS-CoV-2 spike RBD by CD4^+^ T cells. Focusing on the RBD subunit on the spike protein, we have not characterized the full breadth of potential HLA-DR1 epitopes across the spike protein. Similarly, the effect of variant mutations on the breadth of HLA-DR1 epitopes was not fully revealed, as we could focus our in-depth analysis on only a small subset of peptides. Thus, the overall rate of epitope loss within host-viral evolution and whether other mechanisms of escape also exist cannot be determined.

Taken together, the EliteMHCII platform addresses several longstanding challenges in MHC II epitope discovery. It enables direct, scalable, and allele-resolved affinity quantification and supports upstream peptide ranking based on binding strength, reducing false positives and revealing otherwise undetected but biologically relevant epitopes.

Most importantly, EliteMHCII’s integration of biochemical and functional validation creates a uniquely comprehensive system that spans peptide screening to immunogenic confirmation. This end-to-end capacity enhances both fidelity and translational potential, making it a powerful tool not only for basic research but also for clinical applications such as infectious disease monitoring, neoantigen identification, and rational peptide vaccine design. By aligning with population-based benchmarks (e.g., >50% responder frequency in COVID-19 vaccine studies), EliteMHCII advances precision immunology and personalized immune interventions.

## Material and Methods

### RBD peptide pools and biotinylated human CLIP peptide

53 overlapping peptides, each consisting of 15 amino acids with 4 overlapping amino acids, were synthesized from the SARS-CoV-2 RBD protein (aa 319 to aa 541) by GenScript (Nanjing, China) **(Table S4)**. The peptides were dissolved in DMSO at a concentration of 10 mM, stored in a freezer at −80°C, and subsequently diluted in sterile PBS before use. Biotinylated peptides, namely, Bio-CLIP (biotin-Ahx-KPVSKMRMATPLLMQALPM), serve as probes in the detection system also synthesized by GenScript (Nanjing, China). All peptides have a purity exceeding 95%.

### Construction of HLA-DR and HLA-DM expression vectors

The α chain (DRA1*01:01) and β chain (DRB1*07:01, DRB1*03:01, DRB1*15:01, DRB1*11:01, DRB1*01:01, DRB1*13:02, DRB1*13:01, DRB1*04:01, DRB1*11:04, DRB1*04:04, DRB1*15:02, DRB1*09:0, DRB1*14:01, DRB1*01:02, DRB1*15:03, DRB1*12:01, DRB1*04:05, DRB1*10:01, DRB1*04:07, DRB1*04:03, DRB1*12:02, DRB1*16:02, DRB1*04:06, DRB1*14:54) sequences of HLA-DR and HLA-DM (DMA*01:01, DMB*01:03)^41^ were obtained from The IPD-IMGT/HLA Database^42^. We expressed soluble MHC II protein using a knob-into-hole(KIH)-based expression approach. The expression vector employed a heterodimerization strategy based on KIH interactions using human IgG1 Fc^27^. The ectodomains of HLA-DR and HLA-DM alpha chain, featuring Fc with human IgG1Fc CH3 S350C and T362W mutations (konb Fc), along with 6 × histidine tag following alpha chain, were cloned into a pcDNA3.4 modified vector. Simultaneously, the ectodomains of DRB and DMB, incorporating Fc with human IgG1Fc CH3 Y377C, T394S, L369A, Y435V mutations (hole Fc), were cloned into pD2529 vector.

### Recombinant protein expression and purification

All HLA-DR and HLA-DM proteins were expressed in Expi293F cells with OPM-293 CD05 serum-free medium (Cat: 81075-001, OPM Biosciences, China). The α and β chains of HLA-DR and HLA-DM were co-transfected into Expi293F cells using PEI (CAT#24765, Polysciences, USA) transfection reagent and expressed at 37°C for 5 days. For protein purification of HLA-DR and HLA-DM with His-tag, culture supernatant was passed through a Ni-NTA affinity column and further purified by gel filtration (Superdex 200 Increase 10/30 GL, GE Healthcare). The purified HLA-DR protein was stained with uranium formate and observed by negative electron microscopy.

### Affinity Measurement between HLA-DR and CLIP Peptide

For saturation binding, peptide-DR mixtures containing 50 nM HLA-DR, 50 nM HLA-DM, and the biotin-CLIP peptide was started at 100 μM and serially diluted in five-fold, various concentrations of biotin-CLIP were prepared in phosphate buffer saline (PBS) with pH 7.4. HLA-DR was incubated with a biotin-CLIP peptide in the presence of HLA-DM at a 1:1 ratio at 37℃ overnight.

For competition binding, the CLIP peptide was prepared starting at 100 μM and was serially diluted in four-fold. The mixture of CLIP peptide, 50 nM HLA-DR, 50 nM HLA-DM and biotin-CLIP prepared in PBS (pH 7.4) buffer was transferred to a pre-coated 384-well microtitration plate (Maxisop, Nunc, USA) for the capture of peptide-DR by L243 antibody. L243 is a pan-HLA-DR mAb recognizing a conformational epitope in the chain of HLA-DR^43^, The recombinant L243 antibody was expressed from 293F cells and purified using Protein A beads.

After incubation for 60 min at 37°C, followed by washing and reaction with 50 µl Streptavidin HRP (405210, BioLegend). After a further incubation of 30 min at room temperature, followed by five rounds of washing with PBST, 50 µl/well of 3,3′,5,5′-tetramethylethylenediamine solution (SureBlue Reserve TMB Microwell Peroxidase Substrate, KPL) was added and the plates were incubated for 15 min at room temperature. The chromogenic reaction was stopped by the addition of 25 µl 2 M sulfuric acid, and the optical density at 450 nm (OD450) was recorded using a microplate reader (MK3; Thermo Lab system, Helsinki, Finland). Each dilution was repeated in triplicate. The results were interpreted by nonlinear, dose-response regression analysis using GraphPad Prism software. Obtain EC_50_ and IC_50_ affinity values, respectively. The K_d_ values of competitive binding(IC_50_) of CLIP and HLA-DR, converted using the Cheng-Prusoff equation^44^.

### Simulation method

Given that DRB1*1301 and DRB1*1302 proteins differ only at the 170th amino acid position (V170/G170) yet exhibit significant differences in binding energy, and considering their atypical inverted peptide binding orientation, we used molecular dynamics (MD) simulation focused on these two protein systems. We employed AlphaFold3^45^ and HPEPDOCK^46^ docking programs to generate three initial structures of CLIP peptide in complex with DRB1*13:01 and DRB1*13:02 protein, respectively. The protonation state of both six initial structures were predicted by PROPKA^47^, which indicated that H5, H33, D66, H100, E155, and H165 residues at pH = 5.5 becoming protonated, while those residues are neutral at pH = 7.4. Subsequently, MD simulations of the twelve systems (six initial structures at pH = 5.5 and 7.4) were performed using the AMBER24 package with GPU acceleration^48^, and each system ran two parallel simulations.

The AMBER ff19SB force field parameters^49^, TIP3P water model, and 0.15 mol/L NaCl solution setup were employed. Then, MD simulations were performed for each system at 300 K and 1 bar with the isothermal-isobaric ensemble. Each system was equilibrated over 20 ns before the production simulation was run for another 100 ns. All analysis used the last 40 ns trajectory in each system.

### Affinity Measurement between RBD Peptides and HLA-DR

L243 antibody was diluted in 0.05 M NaHCO_3_/Na2CO_3_ buffer (pH 9.6). The wells of high binding 384-well microtitration plates (Maxisorp, Nunc, USA) were coated with 50 µl L243 (100 ng/well) at 4°C for overnight. After incubation, the wells were washed 3 times with PBST and blocked with 100 µl 5% dried silk milk in 0.01 mM PBST (pH 7.4) at room temperature for 1 h. Following three washes with PBST, the plates were dried at room temperature.

The mixture of peptide-MHC II containing 50 nM HLA-DR, 50 nM HLA-DM, 25 nM biotin-CLIP, and competing RBD peptides was serially diluted in PBS buffer at four-fold from 10 μM to 0.6 nM, 37°C overnight incubation. The mixture was transferred to a pre-coated ELISA plate for the capture of peptide-loaded DR by L243 antibody. The plates were assayed using the method described above.

### Peptide binding prediction by NetMHCIIpan

The Peptide binding prediction by NetMHCIIpan assay was described previously^50^. The NetMHCIIpan-4.1 algorithm was used to predict the binding of RBD peptides to DRB1 allotypes^51^. NetMHCIIpan prediction algorithms are trained on data sets of *in vitro* binding affinities and mass spectrometry of MHC II-eluted ligands. The eluted ligand mass spectrometry (EL) score indicates the likelihood of binding, whereas the % rank score is normalized to a set of random peptides. EL score is negatively correlated with percentage rank, which refers to the high binding affinity of the peptide.

### Biotinylation of pMHCII-Fc tetramer

The Biotinylation of pMHCII-Fc tetramer assay was described previously^52^. Expressing MHCII proteins with an avi tag (GLNDIFEAQKIEWHE) at the C-terminus of the α chain. The purified protein was biotinylated with the Avi-tag with BirA. Biotinylated MHCII-Fc was run on Superdex 200 Increase 10/300 GL (Cytiva, US). The biotinylated molecules were then loaded with peptide by incubation with a 10-fold molar excess of RBD peptides and CLIP peptide, 0.2% n-octyl-d-glucopyranoside (Sigma-Aldrich) for overnight at room temperature, and MHCII monomers were incubated with fluorochrome-conjugated streptavidin (SA) at a 4:1 pMHCII: SA molar ratio at room temperature for 30 min. Afterward, 25 μM free biotin was added to block any unoccupied SA binding sites and incubated at room temperature for 10 min. Multimers were subsequently centrifuged at 3300 × *g* for 10 min to remove aggregates and the resulting supernatants were pooled appropriately. and stored at 4℃ and used within a week.

### Donor samples

We collected the peripheral blood from two volunteers who received the Ad5-vectored wildtype strain-based COVID-19 vaccine^53^ two weeks ago and six volunteers received XX vaccine. The HLA-DR typing for the volunteer is shown in Table S5. Peripheral Blood Mononuclear Cells (PBMC) were isolated using density gradient centrifugation in SepMate tubes (StemCell Technologies) containing Ficoll-Paque (StemCell). Blood collected from the leukopaks was diluted 1:1 with 1× PBS (Thermo Fisher). Tubes were centrifuged at 1000 × *g* for 20 min with the centrifuge deceleration brake off to prevent disruption of the Ficoll layer. Plasma and PBMCs were then collected in new 50 mL conical tubes and diluted 1:1 with 1× PBS. The tubes were centrifuged at 450 × *g* for 5 min. Isolated PBMCs were then centrifuged at 450 × *g* for 5 min, the supernatant was discarded, and cells were resuspended in RPMI GlutaMAX media (Thermo Fisher) supplemented with 10% (v/v) FBS.

### Epitope-specific CD4^+^ T cell responses from vaccinees

The Epitope-specific CD4^+^ T cell responses assay was described previously^54^. For in vitro expansion, Fresh PBMCs were in prewarmed RPMI-1640 medium (R0883, Sigma-Aldrich) containing 10% fetal bovine serum (FBS 12-A, Capricorn), 10 mM Hepes (Sigma-Aldrich), 2 mM GlutaMAX (Gibco), penicillin-streptomycin (50 IU/ml; Sigma-Aldrich), and 50 IU interleukin-2 (IL-2; Peprotech) at a concentration of 1 × 10^6^ cells/ml. PBMCs were pulsed with peptides (4 μg/ml) and cultured for 10 to 12 days, adding 100 IU of IL-2 on day 5. In vitro, expanded cells were analyzed by cell surface marker staining. PBMCs from donors treated with zombie Green Fixable Viability Dye (BioLegend, Cat: 423111) at 1:200 and Fc Receptor Blocking Solution (BioLegend, Cat: 422302) in FACS buffer (PBS, 2% FBS) for 20 min at 4℃, 50 nM PE-labeled peptide-HLA tetramers were incubated for 30 min at room temperature. Cells were washed in PBS buffer before surface staining for CD4-BV421 (RPA-T4, eBioscience, Cat: 404-0049-41), CD3-APC (SK7, eBioscience, Cat: 17-0036-42)Stained cells were analyzed on a flow cytometer (BD FACSAria™ II) and analyzed with Flowjo v10.9.0 (BD).

### Immunization of immune system humanized mice

NOD Prkdc^sicd^IL2Rγc^-/-^(NCG) hosts were obtained from GemPharmatech Co. Ltd. (T001475), and housed in specific pathogen-free (SPF) mouse facilities in the Model Animal Research Center, Medical School of Nanjing University. Immune system humanized mice were generated as previously described ^55^. Briefly, the human CD34 MicroBead Kit (Miltenyi Biotec, Cat: 130-046-702) was used for hematopoietic stem and progenitor cells (HSPCs) purification from human fetal liver under clinical ethical approval from the ethical committee of Drum Tower Hospital with informed consent (protocol #2021-488-01). Pups of NCG mice were sublethally irradiated and intrahepatic injected with human CD34^+^HSPCs within 1 week after birth. 12-week-old humanized mice were injected with 50 μg pcDNA3.1(+) vector plasmid containing GM-CSF and IL-4 resuspended in 1.8 mL PBS as previously reported^36^. Four days later, mice were intraperitoneally immunized with a total of 400 μg pooled peptide mixed with 20 μg CpG (GenScript, China). Specifically, we have a high-affinity group and a low-affinity group, and each group has one mouse. High-affinity group received 100 μg of each of four high-affinity peptides, including RBD_323-337_, RBD_407-421_, and RBD_511-525_, whereas the low-affinity group received 100 μg of each of four low-affinity peptides, including RBD_319-333_, RBD_415-429_, and RBD_475-489_. Additionally, a total of 20 μg 4RBD-Fc plus 100 μL Alum adjuvant (InvivoGen, Cat: vac-as03-10, France) were intraperitoneally injected 3 days after peptide immunization. Two weeks later, a booster peptide vaccination was administered with the same formula as the first dose of the vaccine.

### Flow cytometry

Splenocytes of humanized mice were harvested three weeks after the first immunization. Briefly, mice were euthanized, the spleens were taken out and mechanically dissociated, and passed through the 70 μm cell strainer (Corning, Cat: 352350) to obtain single-cell suspension. Then, RBC lysis was performed by treating with ammonium-chloride-potassium (ACK) for 3 min at room temperature. The peptide-HLA-specific cells were identified using HLA-DRB1*01:01 and HLA-DRB1*14:54-restricted tetramers loaded with high-affinity and low-affinity peptides, respectively. The following Abs were used for flow cytometry staining: anti-mouse CD45 (REA737, Miltenyi Biotec, Cat: 130-110-665), anti-human CD45 (QA17A19, BioLegend, Cat: 393408), anti-human CD3 (HIT3a, BioLegend, Cat: 300318), anti-human CD4 (OKT4, BioLegend, Cat: 317416), anti-human CD8α (RPA-T8, BD, Cat: 563795), anti-human CD25 (BC96, BioLegend, Cat: 302636), anti-human CD69 (FN50, BioLegend, Cat: 310910). mAbs staining was performed for 30 min on ice. For *ex vivo* stimulation, splenocytes were incubated with 50 μg/mL corresponding peptide for 2 h, then incubated for an additional 4 h at 37 °C in the presence of brefeldin A (BD). After incubation, cells were washed and stained with mAbs mix. Dead cells were excluded by use of Fixable Viability Dye eFluor 506 (eBiosciences/ Thermo Fisher Scientific). Data were collected on a NovoCyte flow cytometer (Agilent) and were further analyzed with Flowjo v10.9.0 (BD).

### ELISA Detection of RBD-Specific IgM Antibodies in Humanized Mice

Using ELISA to assess the RBD-specific IgM antibody response in humanized mice on the 20^th^ day post-immunization. The 96-well ELISA plates were coated with RBD-His at a concentration of 2 µg/mL in 100 µL per well and incubated overnight at 4°C. Serum samples, serially diluted 100 µL per well, were added and incubated at 37°C for 1 hour. Goat anti-Human IgM Secondary Antibody, HRP (Invitrogen, Cat: 31415), was used to detect the antibodies with an incubation period of 30 minutes at room temperature (100 µL per well). Finally, TMB (3,3’,5,5’-Tetramethylbenzidine) was added at 100 μL per well and incubated at room temperature for 5 minutes, followed by termination with 1M H_2_SO_4_. Washing steps were performed between each step, and absorbance at 450 nm was measured.

### Statistical analysis

By utilizing population coverage frequencies of HLA-DR alleles and the peptide’s high(less than 100 nM), medium (between 100 nM and 1 μM), and low affinities (between 1 μM and 10 μM^22^) for different HLA-DR alleles, the frequencies of HLA-DR populations corresponding to each affinity category were aggregated to determine the overall affinity of the peptide for HLA-DR alleles across the entire population. The correlation between two continuous variables was analyzed using the Spearman correlation analysis. *P* < 0.05 was considered as statistically significant. \**P* < 0.05; \*\**P* < 0.01; \*\*\**P* < 0.001; \*\*\*\**P* < 0.0001; and ns, no significant difference. GraphPad Prism software program version 8.0 was used for data analysis. less than 100 nM as high affinity, between 100 nM and 1 μM as moderate affinity, and between 1 μM and 10 μM.

## Author contributions

Jing Chen contributed to methodology development, validation of results through replication and confirmation, formal analysis, and writing of the original draft. Shang Wu and Ting Zhou were involved in formal analysis and validation of experimental results. Xu Zhu, Chunyu Cheng, and Yan Li contributed to the evaluation of peptide vaccine immunogenicity in humanized mouse models, drafted the initial manuscript section on molecular dynamics simulations, and performed formal analysis. Jun Huo and Hao Dong contributed to drafting relevant sections of the manuscript and conducting formal data analysis. Yuxin Chen was responsible for reviewing and editing the manuscript, including language refinement, structural revisions, and content improvements, and also acquired funding. Xianchi Dong conceptualized the study, contributed to manuscript review and editing, supervised the entire research process, provided essential materials, reagents, and resources, coordinated project administration, and secured funding.

## Expanded View Figure legends

**Figure S1.** Expression of soluble MHC II protein.

(A) Gene construct used for expression of the knob-into-hole (KIH) based MHCII heterodimer.

(B) Representative protein purification of HLA-DRB1*15:01/DRA*01:01 heterodimer. Gel filtration chromatography and SDS-PAGE analysis of HLA-DRB1*15:01/DRA*01:01, which was run under non-reduced condition without βME (left lane) and under reduced condition with βME (right lane). MW, molecular weight markers. βME, beta-mercaptoethanol.

(C) Representative electron micrographs of negatively stained KIH-based MHCII heterodimer.

(D) Schematic overreview of EliteMHCII assay, the saturative binding of MHC-II molecules and antigenic peptides. The detection system includes catalysis, stabilization of MHC-II conformation, peptide editing by HLA-DM, and detection of target biotinylated antigenic peptides, enabling rapid affinity detection for various MHC-II and antigenic peptides.

**Table S1.** Binding free energy (ΔG_bind_, in kcal/mol) and interaction energy decomposition of CLIP peptides with the proteins DRB1*13:01 as well as DRB1*13:02, respectively, under different pH conditions (pH 5.5 and 7.4).

| Protein | pH | $\Delta G_{\text{bind}}$ | | $\Delta E_{\text{vdW}}$ | | $\Delta E_{\text{ele}}$ | | $\Delta G_{\text{pol}}$ | | $\Delta G_{\text{np}}$ | | $\Delta G_{\text{disper}}$ | |
| --- | --- | --- | --- | --- | --- | --- | --- | --- | --- | --- | --- | --- | --- |
|  |  | mean | std | mean | std | mean | std | mean | std | mean | std | mean | std |
| DRB1*13:01 | 5.5 | -39.88 | 9.69 | -113.23 | 7.47 | -865.91 | 37.96 | 868.04 | 36.62 | -92.32 | 4.94 | 163.54 | 7.47 |
| DRB1*13:01 | 7.4 | -39.94 | 10.62 | -108.42 | 6.71 | -1173.01 | 42.12 | 1173.53 | 41.52 | -87.69 | 4.26 | 155.65 | 5.79 |
| DRB1*13:02 | 5.5 | -25.49 | 10.37 | -134.76 | 7.00 | -768.73 | 52.49 | 802.75 | 48.88 | -100.33 | 3.65 | 175.58 | 4.53 |
| DRB1*13:02 | 7.4 | -38.24 | 12.72 | -127.84 | 6.98 | -1089.50 | 43.51 | 1110.67 | 43.77 | -100.30 | 3.29 | 168.72 | 4.35 |

**Figure S2.** Violin plots showing the distribution of Voronoi volumes of hydrophobic core region calculated for the DRB1*13:01 and DRB1*13:02 proteins at different pH conditions (5.5 and 7.4). The white dots represent the average volume and the violin plots include box plots. The line in the box plot represents the interquartile range (IQR). The distribution and density of (A) the entire hydrophobic core region are shown in the left panel, (B) individual residues forming the hydrophobic core are shown in the right panel.

**Figure S3.** Two-dimensional KDE plots representing the distribution of the geometric center of sidechain distances between V170(G170)-F173 and F48-F173 residue pairs in the DRB1*13:01 and DRB1*13:02 proteins at different pH conditions (5.5 and 7.4). F48 and F173 use the sidechain phenyl rings, G170 uses sidechain H atom, V170 uses the carbon atom of the two CH3 groups and the shortest distance between V170 and F173 is selected. (A) DRB1*13:01 at pH 5.5, (B) DRB1*13:01 at pH 7.4, (C) DRB1*13:02 at pH 5.5, (D) DRB1*13:02 at pH 7.4. The colorbar represents the density range from low to high.

**Figure S4.** Distribution of the angles between the N atom of the peptide Q15 (Q15’) sidechain, the Cα atom of the peptide Q15, and the Cα atom of V42 in the DRB1*13:01 and DRB1*13:02 protein systems at different pH conditions (5.5 and 7.4). (A) The distribution of N(Q15’) – Cα(Q15’) – Cα(V42) angles. (B) Representative snapshot of DRB1*13:01 and DRB1*13:02 protein at pH 7.4.

**Figure S5.** The distributions of distance between the nearest heavy atoms of sidechain for different residue pairs in DRB1*13:01 and DRB1*13:02 at different pH values (5.5 and 7.4). The heavy atoms are nitrogen atoms in the sidechains of Arg, Lys, and His residues, and oxygen atoms in the sidechains of Asp and Glu residues. The residue pairs are (A) Peptide R7 (R7’) - E11, (B) Peptide R7 (R7’) - D112, (C) Peptide R7 (R7’) - E155, (D) Peptide K5 (K5’) - E93, (E) Peptide K5 (K5’) - D141, (F) R76 - D141, (G) H5 - D27, (H) D27 - R178 and (I) D112 - E155.

**Table S2.**
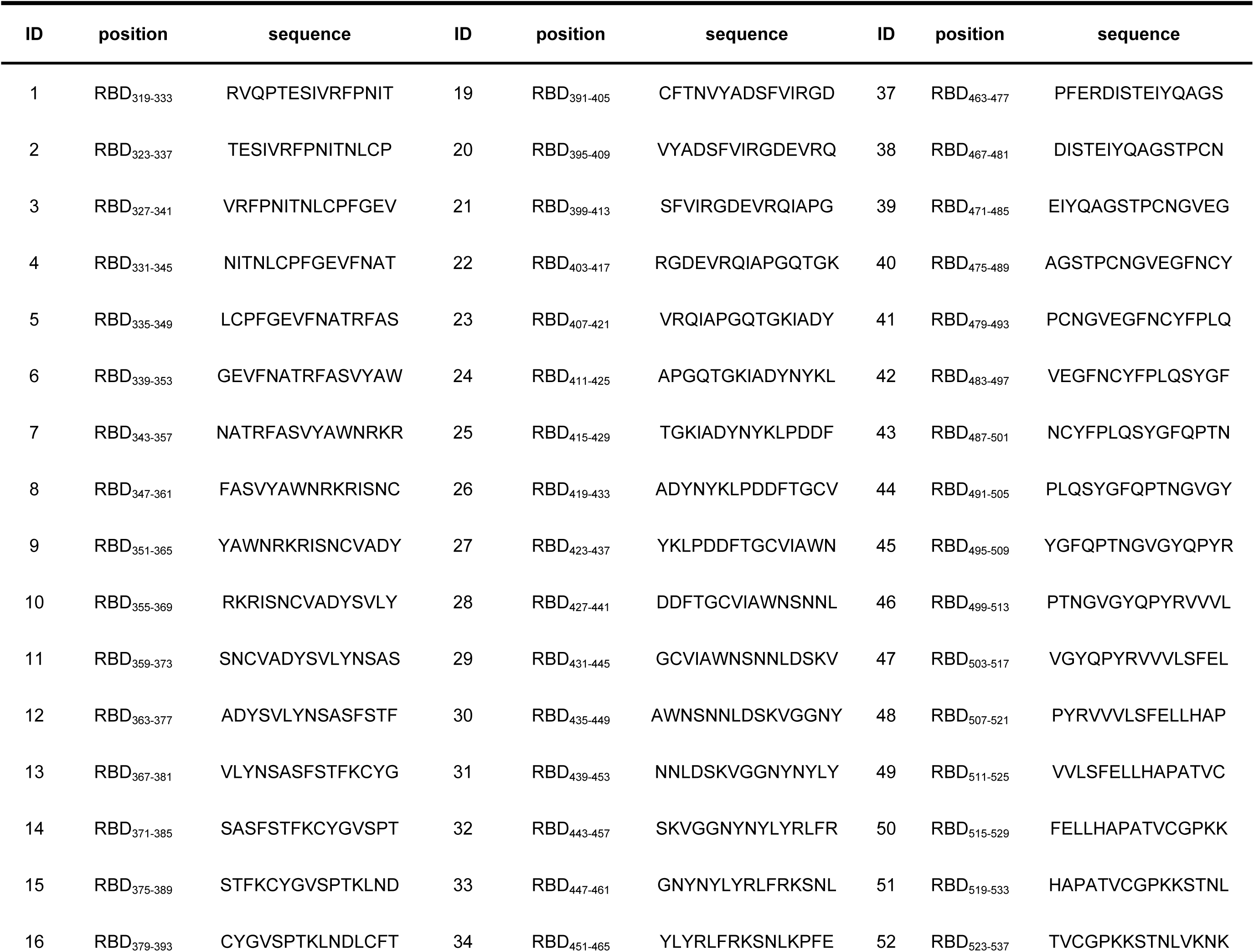
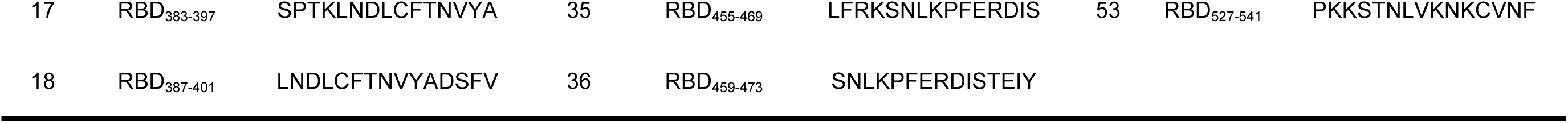
The amino acid sequence of 53 overlapping peptides across RBD protein of SARS-CoV-2 wildtype strain.

**Table S3.** HLA-DR alleles of the volunteers.

| ID | HLA-DR allele |  |
| --- | --- | --- |
| Donor1 | DRB1*07:01 | DRB1*12:01 |
| Donor2 | DRB1*15:01 | DRB1*15:02 |
| Donor3 | DRB1*09:01 | DRB1*07:01 |
| Donor4 | DRB1*09:01 | DRB1*12:02 |
| Donor5 | DRB1*15:01 | DRB1*10:01 |
| Donor6 | DRB1*15:01 | DRB1*11:01 |
| Donor7 | DRB1*15:01 | DRB1*09:01 |
| Donor8 | DRB1*03:01 | DRB1*15:02 |

**Figure S6.** The competitive inhibition assay showing the binding affinity of SARS-CoV-2 RBD_447-461_ derived from wildtype, BA.5, BA.2.86 and JN.1 with 24 popular HLA-DR alleles. The affinity of the RBD_447-461_ epitope from BA.5, BA.2.86 and JN.1 with 12 indicated HLA-DR alleles were reduced compared to that of the wild type (A), whereas their binding affinities with the indicated 12 HLA-DR alleles were comparable with that of wildtype strain (B).

**Figure S7.** The 10 selected peptides were classified based on their binding affinities to 9 HLA-DR alleles from 8 volunteers. Blue indicates high and medium affinity (below 1 µM), orange represents low affinity (between 1 µM and 10 µM), and yellow denotes non-binding peptides (affinity greater than 10 µM).

**Figure S8.** Gating strategy for MHC II tetramer staining of CD4^+^ T cell responses.

(A) Gating strategy to identify MHC II tetramer-specific CD4^+^ T cell responses.

(B) Gating strategy to determine MHC II tetramer-specific CD4^+^ T cells and the AIM^+^ subpopulation.

